# Associations of genetic and spectral diversity in European beech

**DOI:** 10.1101/2023.08.16.553487

**Authors:** Ewa A. Czyż, Bernhard Schmid, Maarten B. Eppinga, Marylaure de La Harpe, Aboubakr Moradi, Cheng Li, Domitille Coq—Etchegaray, Michael E. Schaepman, Meredith C. Schuman

## Abstract

Biodiversity loss presents a growing threat to the global environment, and systematic and spatially contiguous monitoring is needed to inform mitigation strategies. Monitoring of genetic diversity within species, a key factor when assessing biodiversity loss, is laborious and could be supported by scalable phenotypic observations allowing inferences about genetic variation. We studied genetic and phenotypic variation in one of Europe’s most prevalent forest-forming trees, the common beech *Fagus sylvatica* L., using whole-genome sequence data and spectral phenotypes from 219 individuals at 23 sites across the species’ natural range. Spectral phenotypes were collected under standardized illumination and observation conditions from the same top-of-canopy leaves used for nuclear DNA extraction. We found that spectral and environmental information accounted for 77% of the variance along the first two principal coordinates representing genetic structure among sampled individuals, where spectral phenotypes contributed 12% to the prediction of genetic structure. Further, we identified 14 SNPs (single nucleotide polymorphisms), of which two were located within annotated genes, that showed significant associations with variation in leaf reflectance. Our study demonstrates how linking spectral and genomic variation in tree species may be upscaled to the remote observations to support monitoring, understanding and mitigating loss of genetic diversity within species.

## 1. Introduction

Biodiversity loss presents a large and growing risk for the global environment and its ability to support life. Plant diversity loss can negatively affect ecosystem functioning (Purvis et al. 2000, Hector et al. 2007, Isbell et al. 2011, Cardinale et al. 2012, Reich et al. 2012), impact ecosystem stability (Tilman et al. 1994, Hector et al. 2010, Isbell et al. 2015, van Moorsel et al. 2021), and reduce the potential of populations and communities to adapt to environmental change (Des Roches et al. 2018). Biodiversity is under pressure from human activity (Johnson et al. 2009) and global change (Exposito-Alonso et al. 2022). To reduce human impact on biodiversity decline, targets have been set several times aiming to mitigate biodiversity loss worldwide (IPBES 2019), but many further steps need to be taken (Mora et al. 2006, Jenkins and Joppa 2009). One important step towards effective biodiversity maintenance is its continuous and standardized monitoring (Scholes et al. 2008, Turner 2014). Monitoring indicators of biodiversity should help to set conservation priorities (Scott et al. 1989), evaluate actions taken, and develop adaptive management (Sutherland et al. 2004) and policy adjustments to preserve and restore biodiversity (Perrings et al. 2011). Monitoring schemes should span levels of biological organization from ecosystems to species and their genetic diversity. Changes in intraspecific genetic diversity constitute a key component of biodiversity dynamics (Wilson 1988, Exposit-Alonso et al. 2022), as genetic variation provides valuable information about a species’ contribution to ecosystem functioning (Fridley and Grime 2010, Tang et al. 2022) and its evolutionary potential. However, intraspecific genetic diversity is especially challenging to monitor.

The assessment of genetic diversity in the field is based on sample collection, often restricting spatial coverage to a small part of species’ natural range and sampling intensity to a few individuals within a site. Furthermore, genotyping is still commonly limited to small portions of whole genomes, which may not account for all functionally important genetic variation (Lou et al. 2021). Incomplete and biased data inhibits understanding of how species’ phylogeographic history constrains current patterns of genetic variation. Further, these scale-induced limitations result in sparse and biased representations of genetic diversity, which makes it difficult to assess the total adaptative potential of a species to environmental change (Hughes et al. 2008, Des Roches et al. 2018). Whole-genome sequencing to avoid the bias of more restricted sets of molecular markers, linked with acquisitions of phenotypic variation not restricted by spatial extent, may contribute to indicating and understanding species’ genetic diversity under natural conditions.

To collect phenotypic data potentially informative about genetic diversity, imaging spectrometers can provide repeated and spatially contiguous acquisitions of spectral reflectancance of electromagnetic radiation from tree canopies representing integrated signatures of plant biochemical and structural traits (Guyot et al. 1992, Jacquemont and Ustin 2009, Schaepman et al. 2009, Schaepman et al. 2015). These plant traits impact how leaves reflect, absorb, and transmit electromagnetic radiation of different wavelength and thus can affect the spectral phenotypes of individuals from leaf to canopy levels (Meireles et al. 2020). Leaf spectral phenotypes are commonly estimated from spectroradiometer measurements under standardized illumination and observation conditions, while the spectral phenotypes of entire canopies at landscape-scale are derived from remote acquisitions of reflected solar electromagnetic radiation by airborne and spaceborne imaging spectrometers (Schaepman et al. 2009). Because the plant traits shaping spectral phenotypes result from the interaction of plant genes with the environment, spectral acquisitions may facilitate monitoring of genetic diversity under natural conditions, if a link between spectral and genetic variation can be established (Cavender-Bares et al. 2016, Yamasaki et al. 2017). Previous studies associating genetic and spectroscopy data focused on experimental settings (Matsuda et al. 2012, Cavender-Bares et al. 2016, Li et al. 2023) or single-site observational studies under natural conditions, with few within-species assessments and not covering multiple sites across a species’ entire natural range (Madricht et al. 2014, Czyż et al. 2020, Czyż et al. 2023). Furthermore, previous studies were mostly based on genetic markers only covering a small fraction of the species’ genome rather than on whole-genome sequence information.

Here, we address the potential of spectral phenotypes to contribute to assessing intraspecific genetic diversity using short-read whole-genome sequences from 219 individual trees of *Fagus sylvatica* L. (European beech) sampled from across the species range. We chose this species because it is a dominant, ecologically and economically important tree species in Europe that has gained interest as a model tree species for ecological genomics studies under global change (Capblancq et al. 2020). We combined genomic and spectral sampling across 23 sites of *F. sylvatica* using reference-based single nucleotide polymorphism (SNP)-calling to derive genetic diversity, and top-of-canopy leaf reflectance acquired by a contact probe with standard illumination and a consistent background to derive spectral phenotypes. We then use spectral phenotypes, in combination with environmental variables, to predict the genetic structure of the species and to find direct associations of spectra with genetic variation. We link genomics and spectroradiometric approaches with the aim to (1) contribute to current knowledge about genetic diversity in *F. sylvatica* across its natural range; and (2) indicate how spectral phenotypes acquired in natural environments may contribute to prediction of the genetic variation on a continental scale. The presented linkage of spectral with genomic variation from across the species’ natural range indicates how remote observations may be used to support monitoring of genetic diversity in canopy tree species over contiguous areas and on a larger scale than typically attainable when solely relying on field sampling.

## 2. Materials and Methods

### 2.1. Study sites and sample collection

We selected 23 sites dominated by *Fagus sylvatica* L. (common beech) across its natural range in Europe (Fig.1a). We assigned three-letter site names with the first two letters referring to the country that the site is located in, and the third letter referring to the particular location in the country. Most of the sites are covered by protected, old-growth beech forests. The variation among the sites accounts for climatic (mean 50% of the maximum span of environmental values across the range and 95% of the distribution of environmental values across the range), edaphic, and phylogeographic variation (Caudullo et al. 2017). Data were collected *in situ* between 8 August and 25 September 2020, and on 31 July 2021 at the site ROS (Supporting information Tab. S1). At each site we sampled 10 *F. sylvatica* trees (with four exceptions—see following text) that occupied canopy space along two ca. 10 x 200 m transects separated by 50 m. Each transect comprised 5 canopy-forming dominant trees separated by at least 30 m. At some sites we sampled fewer than 10 trees due to a lack of *F. sylvatica* trees in the canopy within the transect (5 trees at RSB, 8 trees at ESP, 7 trees at ROS) and in some cases the sampled material was of insufficient quality for the DNA extraction (1 tree at PLM). When possible, we selected transects on south-west to south-east-facing slopes to minimize variation in solar radiation exposure. From each tree, we collected a top-of-canopy branch of 20 to 100 cm length with the help of tree climbers or, when tree height did not exceed 15 m, with a telescoping carbon fiber pruner (Takeni Trading Co., Osaka, Japan). From each branch, we randomly selected three undamaged leaves (visual inspection) separated by at least 5 cm, from which we acquired spectral data after harvesting the branch (see Materials and Methods, Spectral data) and which we subsequently stored on silica gel for later genetic analysis (see Materials and Methods, Genetic data).

**Fig. 1.**
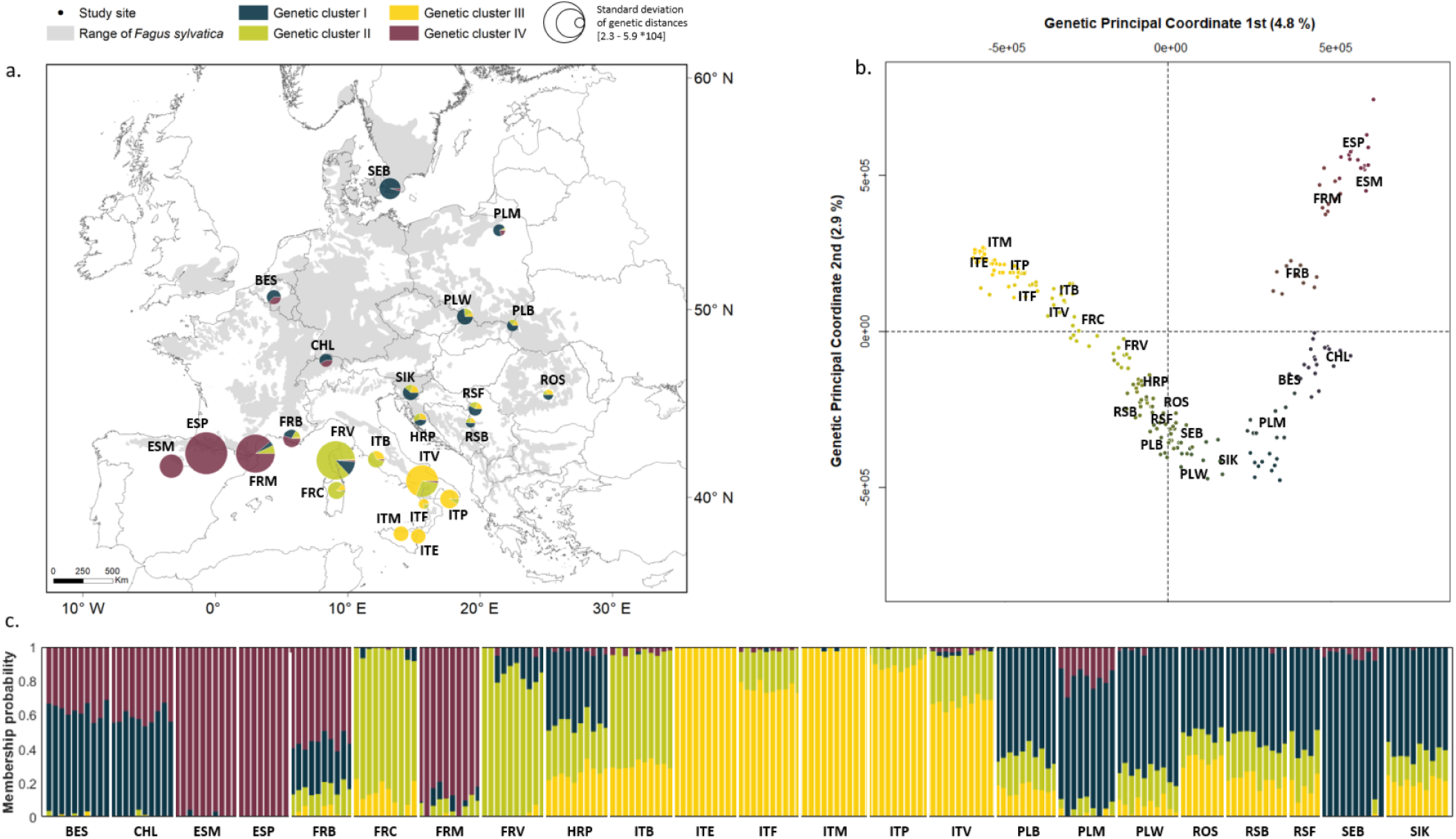
Genetic structure among 23 F. sylvatica sites presented as kinship probabilities of belonging to K=4 genetic clusters with the smallest cross-validation error in ADMIXTURE inferred from 1,380,310 SNPs across the nuclear genome (see Materials and Methods, Genetic data). (a) Averaged kinship probability and the standard deviation (indicated by the circle size) of genetic variation among the individuals from each sampled site (N=10 except for RSB, N=5, ESP, N=8, PLM, N = 9, and ROS, N=7). (b) Relationship between F. sylvatica individuals plotted along the first and second Principal Coordinates (PCOs) derived from genetic distances among individuals. The horseshoe-like distribution of the individuals along the first and second PCOs does not affect the conclusions of the presented analysis (Podani and Miklós 2002) and is consistent with a strongly geographically correlated horseshoe pattern identified in a recent independent analysis of genetic diversity in F. sylvatica (Lazic et al. 2024). Individuals are colored by combined kinship probabilities to four genetic clusters. (c) The membership probability of the 219 sampled F. sylvatica individuals for the derived genetic clusters.

### 2.2. Genetic data

The dried leaves were taken back to the laboratory where we sampled 50 mg of undamaged leaf lamina and stored this in tubes at –20 °C until grinding and DNA extraction. The DNA was purified using a Norgen Plant and Fungi genomic DNA extraction kit (Norgen Biotek, Thorold, ON, Canada), modified to reduce RNA contamination and to improve lysis, yield, and purity. Specifically, samples were flash-frozen in liquid nitrogen immediately prior to grinding and ground using 3 mm glass beads at 30 Hz for 3 min in a TissueLyser II (Qiagen) to a fine powder. The powder was briefly centrifuged before adding lysis buffer and vortexed to mix with the buffer. One μl of RNAse A (DNAse-free, 100,000 units/ml in 50% glycerol, 10 mM Tris-HCl, pH 8.0) and 3 μl Proteinase K (>600 μl ∼ 20 mg/ml in 10 mM Tris-HCl pH 7.5, containing calcium acetate and 50% v/v glycerol) were added to the lysis buffer prior to 10 min incubation at 56°C with 1000 rpm agitation. An additional 1 μl of RNAse A was subsequently added to the reaction, followed by a 10 min incubation at 65°C with 2000 rpm agitation. The samples were then processed according to the manufacturer’s protocol with the modification that 70% ethanol was mixed with lysate by pipetting ca. 10x/sample instead of vortexing. The DNA from clarified lysate was captured on spin columns (centrifugation until all lysate passed) and columns were washed three times with Solution A prior to elution with Elution Buffer B (100 μl, 5 min RT incubation, spinning until all elution buffer was recovered).

The purified DNA was used for sequencing library preparation (Novogene, Cambridge, UK): genomic DNA was randomly sheared into short fragments, which were end repaired, A-tailed, and ligated with Illumina adapters. The fragments with adapters were PCR amplified, size selected, and purified. Libraries were sequenced by Illumina NovaSeq PE150 (Novogene, Cambridge, UK) to produce ca. 11 Gbp/sample, resulting in a mean ca. 20x coverage of the 541 Mbp haploid nuclear genome (Mishra et al. 2022).

Adapters from Illumina were removed using *cutadapt v1.18*. The Illumina reads were trimmed and reads with quality (Q) scores below 30 were removed using *sickle v1.33*. We aligned reads against the chromosome-level reference genome (Mishra et al. 2022) using *bwa-mem2 v2.2.1*. Duplicated reads were marked using *GATK v4.2.4.0* and overlapping read pairs were clipped using *BamUtils v1.0.15*. Mpileup files of all individuals were created using *samtools v1.9* and converted into pro files using *sam2pro v0.8*. From the pro files, nucleotide read quartets at each position of the reference genome were created. Bi-allelic SNPs were detected using a genotype-frequency estimator (Maruki & Lynch 2015) and genotypes were called using the Bayesian genotype caller (Maruki & Lynch 2017) for 32,552,787 SNPs. SNPs with a minor allele frequency below 0.05 and more than 10 missing genotypes across the samples were removed, resulting in 5,252,715 SNPs across all sampled individuals for further analyses. SNPs were pruned for linkage disequilibrium by removing those with a pairwise correlation coefficient (r^2^) > 0.2 within a sliding window of 2 kb using *plink 1.9*, resulting in 1,380,310 SNPs mapped to all 12 beech chromosomes.

### 2.3. Spectral data

Spectral data comprised leaf reflectance spectra collected in the wavelength range from 350–2500 nm using an ASD FieldSpec 4 spectroradiometer (serial nr 18739, ASD Inc., Boulder, CO, USA), coupled to a plant probe with leaf clip attachment, and equipped with a standardized halogen light source and a set of standard black and white reference backgrounds (serial nr 445, ASD Inc., Boulder, CO, USA). Measurements of three leaves from one harvested branch per tree were acquired within minutes to up to 3 h (for ITE, ITM, 5 trees at ITV, and 5 trees at PLB) from the time of branch harvest using the contact probe positioned in the middle of the leaf and to the right of the midrib as viewed from the adaxial side. Measurements comprised five successive readings per sample for four types of acquisitions: a white reference R_w_, a target leaf with white reference as background T_w_, a black reference R_b_, and the target leaf with black reference as background T_b_ (eq. 1). For each reading, five scans, each with an integration time of 8.5 ms, were recorded and averaged per leaf. The averaged reflectance spectra per tree was calculated from the mean of leaf reflectance R using a formula for background-differentiated measurements (Miller et al. 1992):

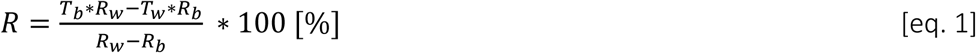

The ASD spectroradiometer provides sampling intervals of 1 nm spectral sampling, although the spectral resolution of the instrument ranges from 3–10 nm. To minimize collinearity effects of neighboring spectral bands and improve the efficiency of analysis, we resampled the spectral signal from the 2151 spectral bands provided by the ASD instrument to 216 spectral bands by linearly interpolating the original signal using the *interp1* function in *Matlab* (v. R2017b) followed by multiplication with a spectral response function (SRF) of Gaussian shape with full width at half maximum (FWHM) of 10 nm, which is the largest nominal FWHM provided in the ASD specifications. These 216 spectral variables were used in all subsequent statistical analyses. For the genome-wide association study (GWAS; see Materials and Methods, Genome-wide association study), we additionally calculated reflectance spectra principal components (PCs) using the *pca* function of *Matlab* (v. R2017b).

### 2.4. Climatic data

Climatic data comprised 19 bioclimatic variables from WorldClim Version2 (Fick et al. 2017) at a resolution of 1 km with global coverage. The correlated variables represent climatic conditions and their annual trends that affect ecological processes across large gradients within the range of *F. sylvatica* and are from the years 1970-2000, thus representing conditions at the time in which the study trees may have been establishing or recently established, rather than the most recent growing seasons. We extracted the value of each environmental variable per site based on the geographic locations of sampled sites and applied min-max normalization of extracted values for each variable. Note that all trees at a given site were thus assigned the same values for the climatic covariates.

### 2.5. Statistical analysis

#### 2.5.1. Analysis of genetic structure

First, we used the filtered SNPs (see Materials and Methods, Genetic data) to estimate genetic structure and relatedness of individuals. Genetic structure was calculated in *ADMIXTURE* v. 1.3 (Alexander et al. 2015) and the number of genetic clusters was set to 4 based on the smallest cross-validation error (Supporting information Fig. S1). To derive the relationships among trees, we calculated the SNPs-based pairwise distance between individuals using *plink* v. 1.9, and applied the *cmdscale* function of *Matlab* (v. R2017b) to perform principal coordinate analysis (PCoA) and to derive the genetic principal coordinates (PCOs). Based on pairwise distances between individuals, we additionally calculated within-site standard deviations of genetic relatedness.

#### 2.5.2. Redundancy analysis

Furthermore, we estimated independent and common contributions of climatic and spectral information to model genetic structure among sampled *F. sylvatica* sites. We used redundancy analysis (RDA) as a multivariate ordination method to simultaneously account for multiple (genetic) response variables (Capblancq and Forester 2021). We used Akaike Information Criterion (AIC) values for forward selection of (climatic or spectral) explanatory variables and a threshold of 2.5 for Variance Inflation Factor (VIF) scores to reduce the impact of collinearity of these variables (Johnston et al. 2018). We limited the (genetic) response variables to the first and the second genetic PCO. With n = 23 sites to analyze the correlation structure between climatic and genetic variables we aimed to keep the total number of explanatory and response variables low to avoid overfitting (with 22 variables perfect fits would be obtained).

Firstly, we constructed RDA models from climatic and spectral information separately. For the RDA model with climatic data, we retained four climatic explanatory variables with low correlation (out of 19 total), while for the RDA model with spectral data we retained four spectral explanatory variables (out of 216 total) as mutually uncorrelated. We then combined three climatic and four spectral explanatory variables with low correlation (one of the climatic variable was excluded due to the correlation with the spectral variables) with the two genetic PCOs as response variables for a combined RDA model. We partitioned the contributions of each climatic and spectral variable to the total genetic variation (as variance along genetic PCO1 and PCO2) using analysis of variance (ANOVA). We used the *rda* function of the *vegan* package (v. 2.6-4) in *R* (v. 4.1.3) to construct the models and to find the model fits.

#### 2.5.3. Genome-wide association study

In addition to assessing the contribution of climatic and spectral variation to the total variation in the first to genetic PCOs, we also associated the 1,380,310 filtered SNPs across the genome with spectral variation while accounting for climatic differences between sites. For that, we conducted a genome-wide association study (GWAS) using the Bayesian-information and Linkage-disequilibrium Iteratively Nested Keyway (BLINK) model (Huang et al. 2019) implemented in *GAPIT* (v. 3) (Wang and Zhang, 2021) in *R* (v. 4.1.3). We used the first four genetic principal components (as calculated in *GAPIT* and each accounting for more than 1% of total genetic variation) to account on the species’ genetic structure and environmental variables as covariates. The *BLINK* model controls for false negatives by accounting for the linkage disequilibrium of SNPs across the genome and the covariates in the associations. We consecutively derived the proportion of variance in spectral phenotypes explained by genotypic variance for associations that were significant after applying a Bonferroni correction that accounts on the multiple statistical tests to reduce false positives. This task was performend in *GAPIT* software. We further compared the locations of the significantly associated SNPs to the annotated genome of *F. sylvatica* to determine which SNPs were located within annotated genes. For these SNPs, we indicate the corresponding gene ontology (GO) terms as aligned with the annotated reference genome Bhaga v.2 (Mishra et al. 2022).

## 3. Results

### 3.1. Genetic structure of *F. sylvatica* across the species natural range

We present an assessment of genetic structure of a broad-leaved tree species across its natural range derived from whole-genome sequences of nuclear DNA. Using ADMIXTURE cross-validation, we identified four genetic clusters across the current geographic distribution range of *Fagus sylvatica* L. (Fig. 1). We identified a high abundance of one genetic cluster in southern Italy (ITE, ITM, ITP, ITV; cluster indicated in yellow), one genetic cluster in the southern Pyrenees (ESP, ESM, FRM; cluster indicated in maroon), and one genetic cluster centered in Corsica (FRV, FRC; cluster indicated in green). At the other sites individuals could not be clearly assigned to a single cluster, or were collectively assigned to a fourth cluster (indicated in blue), suggesting that except for the above sites located at the extremes of the distribution range beech has weaker genetic structure and rather seems to represent a large panmictic population (Fig. 1a), with an exception of the population at SEB, the closest site to the northern extreme of the species distribution, dominated by the blue cluster.

Using the first two genetic PCOs to represent the genetic distances between individuals and sites supports the finding of a continuous genetic distribution with geographically more extreme populations representing enpoints along a continuum of genetic relatedness (Fig. 1b). The pairwise genetic distances demonstrate that trees within sites are genetically more similar than between sites, while the standard deviations of genetic distances within populations (represented by circle size in Fig. 1a). The sampled sites in central and northern Europe are characterized by a relatively small standard deviation of genetic distances within sites (Fig. 1a). In contrast, per-individual kinship probabilities to different genetic clusters, and pairwise genetic dissimilarities based on allele frequencies between individuals, demonstrate relatively higher within and between-population variation in southern as compared to northern samples (Fig. 1a, b).

### 3.2. Genetic variation explained by climatic and spectral variation

Together, environmental climatic variation between sites and spectral information variation within and between sites explained 77.2% of the total genetic variation represented by the first and second genetic PCO of sampled *F. sylvatica* (Fig. 2, Supporting information Fig. S2). Environmental climatic variables explained 42.6% of the total genetic variation independently of spectral information, *i.e.*, after fitting spectral information first in the model. The environmental climatic variables contributing most were Mean Temperature of Driest Quarter (BIO 09), Isothermality (BIO 03), Precipitation Seasonality (BIO 15) and Precipitation of Driest Quarter (BIO 17). In contrast, the first four PCs obtained from a PCA of the 216 spectral variables explained 12.1% of the total genetic variation among trees additionally to environmental climatic variables, *i.e.*, after fitting climatic information first in the model. This lower percentage must be viewed in the context that climatic variables did not vary within sites, while spectral variables did. Portions of spectra which were most informative about genetic variation corresponded to variation of spectral phenotypes were centered at 350 nm (ultraviolet, UV), 430 nm and 530 nm (visible, VIS), and 1500 nm (shortwave infrared, SWIR). 22.5% of the total genetic variation explained by environmental climatic or spectral variation could not be separated, *i.e.*, was shared by the two (Fig. 2).

**Fig. 2.**
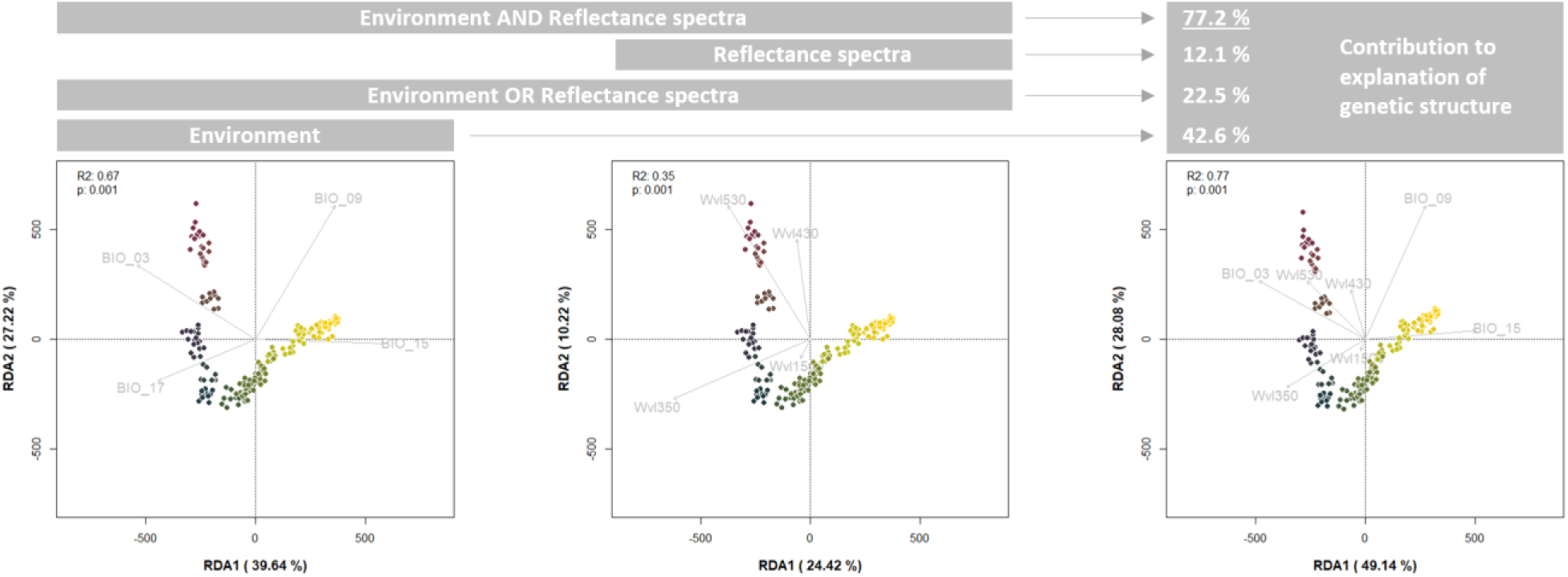
RDA models for prediction of F. sylvatica genetic variation as captured by the first (PCO1) and second (PCO2) genetic Principal Coordinates across the species range derived from environmental climatic and spectral information separately and environmental climatic and spectral information combined. The explanatory variables were chosen based on AIC and VIF <2.5. Out of 19 environmental climatic variables, four were identified to be indicative of the genetic variation between sites and out of 216 spectral bands, four were indicative of the genetic variation between and within sites. The full model combined three out of the four selected environmental climatic variables for the climate RDA with the four spectral variables selected for the spectral RDA. The incorporation of spectral information added 12.1% explanatory power to a model based on environmental climatic information only (42.6% + 22.5% = 65.1%), yielding a total of 77.2% explanation of variation along genetic PCO1 and PCO2 (mean R^2^ for PCOs 1 and 2) among sampled F. sylvatica individuals explained by climatic and spectral information. The F. sylvatica individuals in RDA space are colored by combined kinship probabilities to four genetic clusters.

### 3.3. Associations between genetic and spectral variation

We identified 14 SNPs across the whole genome of which variation was significantly associated with variation of leaf reflectance spectra or their PCs (Fig. 3). Out of 14 recognized SNPs, four were associated with three distinct wavelength ranges across the reflectance spectra, and ten SNPs were associated with eleven unique reflectance spectra PCs. The spectral ranges across which reflectance varied significantly with genetic variation were 360 nm (significantly associated with two SNPs on the 1^st^ and 8^th^ chromosome), 1910–1960 nm (significantly associated with two SNPs on the 2^nd^ and 5^th^ chromosome), and above 2450 nm (significantly associated with the same SNP on the 5^th^ associated with 1910-1960 nm and with an additional SNP on the 2^nd^ chromosome). The proportion of variation in reflectance spectra explained by genetic variation reached 77.89 % for the association of SNP on the 2^nd^ chromosome (position 51570204) with a wavelength at 1950 nm (p-value 2.15×10^-8^) and 39.55 % in variation of reflectance spectra PCs for the association of an SNP on the 9^th^ chromosome (position 3954556) with the 108^th^ reflectance spectra PC (p-value: 4.68×10^-12^). The variation of the 169^th^ reflectance spectra PC (being the highest PC that significantly associated with genetic variationexplaining less than 2.8×10^-8^ in the variation of reflectance spectra) was explained 7.36 % and 10 % by variation of two SNPs located on the 9^th^ chromosome (position 11662665, p-value: 2.61×10^-8^ and position 11661800, p-value: 2.19×10^-8^, respectively).

**Fig. 3.**
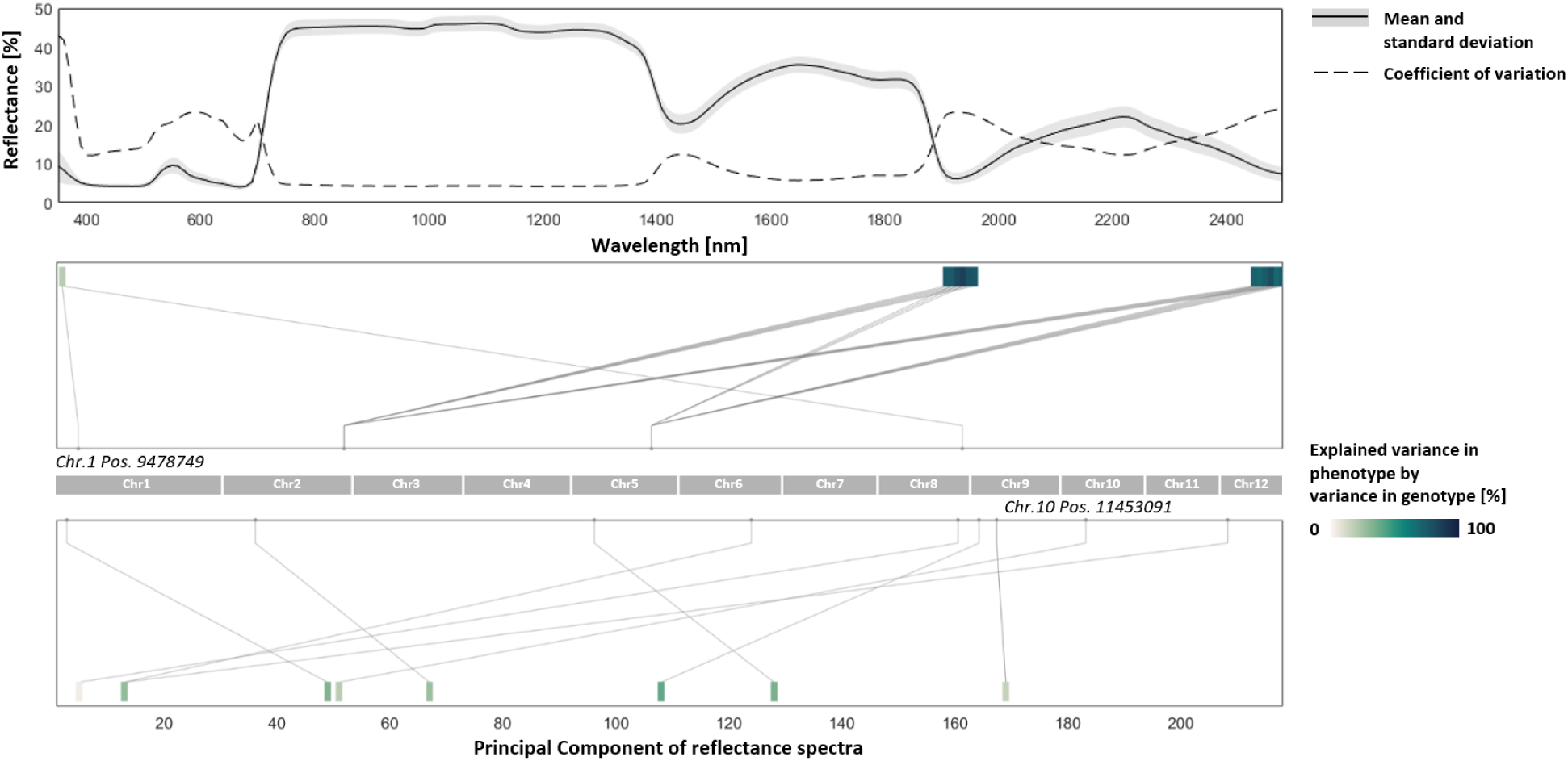
Variation in reflectance spectra (phenotype) and associations of variation in 216 spectral variables and their PCs with variation in genotype as SNPs derived from whole-genome sequences of F. sylvatica. The top panel represents the mean, standard deviation, and coefficient of variation of leaf reflectance spectra from leaves collected from 219 trees. The middle panel represents significant associations between variation across particular wavelengths and four SNPs along the genome. We note that the measurement uncertainty is higher in regions where the coefficient of spectral variation is higher, particularly for wavelengths below 400 nm (Petibon et al. 2021). The bottom panel represents eight spectral principal components significantly associated with variation at ten SNPs across the genome; it should be noted that these are high-dimensional PCs that thus represent a small proportion of the total spectral variation (less than 1%). The shading of the associations indicates the variance explained in a particular phenotype (wavelength or spectral PC) by all associated SNPs. The two labeled loci across the genome correspond to the annotated genes.

Out of 14 SNPs that significantly associated with variation in reflectance spectra or their PCs, we identified two SNPs within annotated genes of *F. sylvatica* (Supporting information Tab. S1). The associated GO terms included plasma membrane, integral component of membrane, magnesium ion binding, ATP binding, hydrolase activity, ATPase-coupled intramembrane lipid transporter activity (for SNP at the position 11453091 on the chromosome 10^th^), and phospholipid and serine-type endopeptidase (for SNP at the position 9478749 on the chromosome 1^st^). The first SNP was associated with a spectral wavelength of 360 nm, and the second was associated with the 51^st^ PC derived from the 216 spectral variables. It is important to note that wavelength 360 nm is in the spectral range of high measurement uncertainty (Petibon et al. 2021) and that the 51^st^ PC describes 9.2 x 10^-6^ % of the total spectral variation.

## 4. Discussion

By merging genomic with spectral approaches we present the potential of spectral phenotypes to contribute to the predictions of intraspecific genetic variation of a forest tree species across its natural range. We note that the contribution of spectral variaton is, in our dataset, the only factor that distinguishes within-site variation, whereas the climatic variables we use are appropriate to distinguish among, but not between sites. Correspondingly, the climatic variables predict a larger proportion of the total genetic distance measured across the species range. The spectral information we collected contributed to the prediction of neutral genetic structure across the sampled populations. We also identified a few genomic variants that are significantly associated with specific components of spectral variation, although some of the associations are within spectral ranges of high measurement uncertainty or with PCs explaining less than 1% of spectral variance. Predictions of spatial patterns in genetic variation may be further improved by incorporation of the species’ phylogeographic history and functional genomics resources, and hold the potential to be further guided by spectral information remotely accessed a landscape scale and repeatedly over time.

### 4.1. Species phylogeographic history

While spectra carry information about current genetic composition, further incorporation of information about the species’ phylogeographic history explaining the development of genetic patterns could help constrain genomic predictions to the available gene pool of particular areas. The genetic clusters indicated in yellow and maroon in Fig. 1 may correspond to previously reported glacial refugia (Magri et al. 2006), while further genetic and phylogeographic studies would be valuable in Corsica to determine whether the cluster indicated in green may represent intermixing between refugia or an additional refugium (Petit et al. 2003). The origin of the most widespread genetic cluster (indicated in blue) in central and northern Europe remains uncertain based on our sampling and might be assigned to the eastern range of the Alps (Gomory et al. 2003) or southern Moravia (Magri et al. 2006). When the most northern site within the species range dominated by the blue cluster may have been a result of a bottle-neck. The phylogenetic structure across the range of *F. sylvatica* is thought to result from post-glacial expansion: from the west across the central European plains (cluster indicated in maroon), from the south across the Carpathian mountain range (cluster indicated in yellow), and from Corsica and nearby sites at the French Mediterranean basin (FRB, FRM) across the Appenninan Peninsula (ITB, ITV, ITF, ITP), Balkans (HRP, RSB, RSF, SIK) and the Carpathian mountain range (PLB, PLW), with the last among these likely permitted by terrestrial connections between the Adriatic and Mediterranean coasts during the last glaciation (cluster indicated in green) (Hazler et al. 1997). These genetic patterns correlate with environmental niches defined by water availability (BIO 09, BIO 15, BIO 17) and isothermality (BIO 03), which may determine adaptive patterns in European beech (Fang and Lechowicz 2006, Bolte et al. 2007), blurred by gene flow. In contrast to lower mountain ranges, the Alps likely act as a barrier for gene flow between southern and northern populations (Paule 1995). Additionally, the continuous distribution of the species in the northern part of its range likely allows greater gene flow across the landscape, while the patchier distribution in the south, with a more restrictive topography an thus climatic variation, may limit gene flow and maintain genetic distance between populations. This may result in greater genetic differences between southern sites, which may have further been enhanced by a longer history and post-glacial climatic changes causing migration to higher altitudes (Thiébaut et al. 1982). Merging the current genetic variation and the phylogeographic history of the species with regularly monitored spectral (phenotypic) variation could allow more accurate assessment of genetic diversity within tree species across its range.

### 4.2. Associations of spectral variation with genetic structure and genetic variants

We report spectral-genetic associations based on spectral phenotyping of leaves sampled at selected sites across the natural range of *F. sylvatica*. These associations showed that the spectral variation that contributed most to the prediction of genetic structure in this study corresponds to pigment absorption features related to electronic transitions induced by light at 410 nm and 530 nm (Gitelson et al. 2006), leaf water content related to vibrational absorption of photons at 1500 nm, and potentially leaf surface morphological properties affecting the reflection of UV light at 350 nm, with the caveat that this last signal is characterized by higher uncertainty (Petibon et al. 2021). This suggests that the species’ genetic diversity may be predicted by these spectral variables as proxies of the underlying leaf traits. This additionally requires that variation in these leaf traits is substantially genetically based and less strongly influenced by environmental variation.

Additonally, we identified loci across the genome significantly associated with reflectance in UV radiation around 360 nm and in SWIR above 1800 nm and for the eleven distinc PCs of reflectance spectra. Among the 14 SNPs showing significant associations with spectral features two were located within annotated genes. One gene (on the 1^st^ chromosome) is annotated as having homology to genes in other organisms that are known or predicted to encode a P-type ATPase associated with magnesium ion transport and ATPase-coupled intramembrane lipid translocation—functions central to maintaining chloroplast structure and ion homeostasis (Dukic et al. 2023; Østerberg et al. 2018). Given magnesium’s essential role as the central atom in chlorophyll, variation in its regulation may influence pigment composition and thereby modulate spectral reflectance in the UV-visible range. Notably, this association showed an exceptionally small p-value (2.59 × 10⁻²⁷), standing out from the rest of the significant associations, but was not supported by nearby SNPs in linkage disequilibrium. This raises the strong possibility that this is a false positive (Hinney et al. 2007) or the result of unaccounted confounding factors (Grinde et al. 2024) potentially impacted by high measurement uncertany in this spectral range (Petibon et al. 2021). The second gene (on the chromosome 10^th^) is annotated as a putative serine-type endopeptidase, a class of enzymes involved in proteolytic processing during development and stress responses (Zhang et al. 2006). Serine proteases have also been implicated in membrane remodeling and lipid-protein interactions (Higashi et al. 2019), processes that can affect leaf surface properties and structural organization that could be relevant to SWIR reflectance. Further studies identifying genetic pathways in which the localized genes are involved would potentially help to understand the mechanistic foundation of genetic drivers of particular plant traits that influence these spectral variations in our model species. In addition, polygenic, structural genomic, and epigenic impacts on spectral variation could further facilitate research on selection processes that drive adaptations in functional genes across broad spatial and temporal scales. Finally, genetic manipulation studies would be required to test causal links in these associations. The use of experimental setups manipulation genotype-by-environment interactions would allow to test associations and potentially to quantify heritability. Studies conducted across networks of common gardens as well as studies manipulating environmental and genetic factors would be required to separate genetic and environmental influences on spectral variation. Common garden networks are increasingly available for *F. sylvatica* and should be used to test spectral–genetic associations.

We acknowledge that the derived spectral–genomic associations are specific to the sampled individuals across the environmental gradient covered by the 23 sites. Furthermore, we note that our analyses did not correct for potential confounding effects of sampling conditions such as weather, daytime, season or for environmental heterogeneity within sites. We generally sampled from southern to northern sites over time in order to sample sites at similar Growing Degree days (GDD) (Supporting information Tab. T1) and sampled transects were within a similar aspect such that environmental variation would be primarily in microenvironmental factors such as localized shading or differences in belowground conditions; sampling was furthermore limited to large canopy-forming trees that should experience more similar microenvironments within a site. Additionally, our analysis includes a widely adopted publicly available climate dataset that can be interpreted as representing climatic conditions earlier in the establishment and growth of the studied trees and is too coarse-grained to capture within-site differences. Thus in our case, it is clear that leaf spectral data include complimentary information about the current state of trees as well as within-site variation among sampled individuals. The inclusion of more current and more spatially resolved environmental information would allow to estimate the genetic contribution to leaf spectral variation as measured within and across these sites. Expansion of the sampling to genetically or environmentally unique sites, and further randomization or standardization of collection across weather, temporal, and microclimatic conditions would additionally expand the scope to predict genetic variation from spatially broad and contignuous spectral information.

### 4.3. Potential of remote observations to contribute to monitoring genetic variation

We have used the largest sampling to date of genomic and spectral variation of *F. sylvatica* across its natural range, combining genetic variation derived from short-read whole-genome sequencing with variation in spectral phenotypes of leaves. We have shown a potential contribution of spectral information to support predictions of genetic diversity. This represents an example how to move on from previous studies carried out at single sites and with restricted genetic information (Czyż et al. 2020, Czyż et al. 2023, Li et al. 2023, Cavender-Bares et al. 2016, Seeley et al. 2023, Meireles et al. 2020) to large-scale remote assessments of genetic variation and presents the potential to associate spectral variation within species with functional genes which may futher complement the remote mapping of plant functional traits. As this study presents a proof of concept for intraspecific genetic variation assessments under natural conditions, it relies on the spatially sparse *in situ* leaf-level investigation of spectral phenotypes and is thus only a first step from which upscaling to remote observations of spectral phenotypes at a landscape scale and beyond will have to be developed.

Spectral phenotypes at the canopy level acquired from remote observations of airborne and spaceborne imaging spectrometers will allow for spatially dense spectral sampling over large geographical areas. Data from existing spaceborne imaging spectrometers, and the wealth of planned and upcoming imaging spectrometer missions (including NASA’s SBG (Cawse-Nicholson et al. 2021) and ESA’s CHIME (Celesti et al. 2022)), will bring spatially contiguous and temporally dense information about spectral properties of the Earth surface resolved at spectral intervals which are sufficiently narrow to be interpreted in terms of molecular absorption features. However, the potential correlation between genomic and spectral variation acquired over large areas and at coarser spatial resolution with pixels each representing a plant community rather than a single individual (Zheng et al. 2022, Helfenstein et al. 2022) and with a lower Signal-to-Noise ratios of spectral signal still need to be evaluated in terms of their potential contribution to monitoring genetic diversity supported by remote observation.

Spectral support of predictions for genetic diversity across space and time may ultimately help to target environmentally, phenotypically, and genetically underrepresented sites or hotspots for the monitoring and conservation of genetic resources and contribute to calibration of predictive models. Improved predictions of genetic variation supported by spectral phenotypes could be used to develop novel indicators of change in genetic diversity and adaptation within tree species over time when combined with repeated monitoring and could deliver information for the adjustment of current efforts to safeguard biodiversity at the global scale, at unprecedented speed and spatial completeness.

## Acknowledgments

This study was supported the University of Zurich Research Priority Program on Global Change and Biodiversity (URPP-GCB), and funded by the NOMIS grant on “Remotely Sensing Ecological Genomics”. We express our gratitude to Emmanuelle Argenti, Ventura Gennaro, Hugot Laetitia, Elodie Magnanou, Joseph Garrigue, Kris Vandekerkhove, Arne Verstraeten, Nilsson Jorgen, Piotr Małż, Katarzyna Gierad, Dejan Stojanović and the Institute of Lowland Forestry and Environment; University of Novi Sad, Gordan Lukač, Katja Konečnik, Janez Šubic, Damen Kocjan, Eustaquio Gil Pelegrín, Juan Pedro Ferrio, Piotr Poloczek and Stanisław Kucharzyk for allowing access to their sampling sites and support on investigation and to Romeo Galiano, Joseph Garrigue, Zbigniew Stebel, Helgessons TrädTjänst AB company and Michał Szydłowski for assistance in sample collection. We express our gratitude to Reinhard Furrer, Anna K. Schweiger, Benjamin Dauphin, Katalin Csilléry, Christian Rellstab and Ingmar Staude for helpful discussions. A portion of this research was carried out at the Jet Propulsion Laboratory, California Institute of Technology, under a contract with the National Aeronautics and Space Administration. © 2025 all rights reserved. Government sponsorship is acknowledged.

## Declaration of Competing Interest

The authors declare that they have no known competing financial interests or personal relationships that could have appeared to influence the work reported in this paper.

## Author contributions

Initials are listed in order of contribution. Conceptualization: EAC, MES, BS, MBE, MH, MCS. Methodology: EAC, MES, MCS, BS, MBE, AM, MH. Software: EAC, BS, CL. Validation: EAC, BS, MBE. Formal analysis: EAC, BS, MBE. Investigation: EAC, BS, MBE, MH, CL, MCS, MES. Resources: MES, MCS. Data Curation: EAC, DLE. Writing - Original Draft: EAC. Writing - Review and Editing: EAC, BS, MBE, MH, MCS, CL, DLE, MES. Visualization: EAC. Supervision: MES, MCS, BS. Project administration: MCS, MES. Funding acquisition: MES.

## Data availability

The raw short-read whole genome sequences genomic data that support the findings in this study are deposited at NCBI Sequence Read Archive (SRA) under the project with accession number PRJNA980459. The leaf spectroscopic data are archived in the Specchio Spectral Information System (https://specchio.ch/). All data will be publicly available upon publication.

## Code availability

The bash codes developed for deriving genetic structure, and *R* and *Matlab* codes developed for the statistical analysis are deposited at GitHub (*to be established upon the publication*) and will be publicly available upon publication.

## Notes

### Competing Interest Statement

The authors have declared no competing interest.

### Summary of Updates

The scope of the manuscript was extended to spectral-genomic associations. Additionally, one extra population of the species of interest was included to the study.

## References

Alexander, D. H., Shringarpure, S. S., Novembre, J., & Lange, K. (2015). Admixture 1.3 software manual. Los Angeles: UCLA Human Genetics Software Distribution.

Bolte, A., Czajkowski, T., & Kompa, T. (2007). The north-eastern distribution range of European beech—a review. Forestry, 80(4), 413–429.

Capblancq, T., & Forester, B. R. (2021). Redundancy analysis: A Swiss Army Knife for landscape genomics. Methods in Ecology and Evolution, 12(12), 2298–2309.

Capblancq, T., Morin, X., Gueguen, M., Renaud, J., Lobreaux, S., & Bazin, E. (2020). Climate-associated genetic variation in Fagus sylvatica and potential responses to climate change in the French Alps. Journal of Evolutionary Biology, 33(6), 783–796.

Cardinale, B. J., Duffy, J. E., Gonzalez, A., Hooper, D. U., Perrings, C., Venail, P., … & Naeem, S. (2012). Biodiversity loss and its impact on humanity. Nature, 486(7401), 59–67.

Caudullo, G., Welk, E., & San-Miguel-Ayanz, J. (2017). Chorological maps for the main European woody species. Data in brief, 12, 662–666.

Cavender-Bares, J., Meireles, J. E., Couture, J. J., Kaproth, M. A., Kingdon, C. C., Singh, A., … & Townsend, P. A. (2016). Associations of leaf spectra with genetic and phylogenetic variation in oaks: prospects for remote detection of biodiversity. Remote Sensing, 8(3), 221.

Cawse-Nicholson, K., Townsend, P. A., Schimel, D., Assiri, A. M., Blake, P. L., Buongiorno, M. F., … & SBG Algorithms Working Group. (2021). NASA’s surface biology and geology designated observable: A perspective on surface imaging algorithms. Remote Sensing of Environment, 257, 112349.

Celesti, M., Rast, M., Adams, J., Boccia, V., Gascon, F., Isola, C., & Nieke, J. (2022, July). The Copernicus Hyperspectral Imaging Mission for the Environment (Chime): Status and Planning. In IGARSS 2022-2022 IEEE International Geoscience and Remote Sensing Symposium (pp. 5011–5014). IEEE.

Csilléry, K., Lalagüe, H., Vendramin, G. G., González-Martínez, S. C., Fady, B., & Oddou-Muratorio, S. (2014). Detecting short spatial scale local adaptation and epistatic selection in climate-related candidate genes in E uropean beech (F agus sylvatica) populations. Molecular Ecology, 23(19), 4696–4708.

Cuervo-Alarcon, L., Arend, M., Müller, M., Sperisen, C., Finkeldey, R., & Krutovsky, K. V. (2021). A candidate gene association analysis identifies SNPs potentially involved in drought tolerance in European beech (Fagus sylvatica L.). Scientific Reports, 11(1), 2386.

Czyż, E. A., Guillén Escribà, C., Wulf, H., Tedder, A., Schuman, M. C., Schneider, F. D., & Schaepman, M. E. (2020). Intraspecific genetic variation of a Fagus sylvatica population in a temperate forest derived from airborne imaging spectroscopy time series. Ecology and evolution, 10(14), 7419–7430.

Czyż, E. A., Schmid, B., Hueni, A., Eppinga, M. B., Schuman, M. C., Schneider, F. D., … & Schaepman, M. E. (2023). Genetic constraints on temporal variation of airborne reflectance spectra and their uncertainties over a temperate forest. Remote Sensing of Environment, 284, 113338.

Des Roches, S., Post, D. M., Turley, N. E., Bailey, J. K., Hendry, A. P., Kinnison, M. T., … & Palkovacs, E. P. (2018). The ecological importance of intraspecific variation. Nature ecology & evolution, 2(1), 57–64.

Dukic, E., Van Maldegem, K. A., Shaikh, K. M., Fukuda, K., Töpel, M., Solymosi, K., … & Spetea, C. (2023). Chloroplast magnesium transporters play essential but differential roles in maintaining magnesium homeostasis. Frontiers in Plant Science, 14, 1221436.

Exposito-Alonso, M., Booker, T. R., Czech, L., Gillespie, L., Hateley, S., Kyriazis, C. C., … & Zess, E. (2022). Genetic diversity loss in the Anthropocene. Science, 377(6613), 1431–1435.

Fang, J., & Lechowicz, M. J. (2006). Climatic limits for the present distribution of beech (Fagus L.) species in the world. Journal of Biogeography, 33(10), 1804–1819.

Fick, S. E., & Hijmans, R. J. (2017). WorldClim 2: new 1-km spatial resolution climate surfaces for global land areas. International journal of climatology, 37(12), 4302–4315.

Fridley, J. D., & Grime, J. P. (2010). Community and ecosystem effects of intraspecific genetic diversity in grassland microcosms of varying species diversity. Ecology, 91(8), 2272–2283.

Gallois, A., Audran, J. C., & Burrus, M. (1998). Assessment of genetic relationships and population discrimination among Fagus sylvatica L. by RAPD. Theoretical and applied genetics, 97, 211–219.

Gausman, H. W., Allen, W. A., Escobar, D. E., Rodriguez, R. R., & Cardenas, R. (1971). Age Effects of Cotton Leaves on Light Reflectance, Transmittance, and Absorptance and on Water Content and Thickness 1. Agronomy Journal, 63(3), 465–469.

Gitelson, A. A., Keydan, G. P., & Merzlyak, M. N. (2006). Three-band model for noninvasive estimation of chlorophyll, carotenoids, and anthocyanin contents in higher plant leaves. Geophysical research letters, 33(11).

Gomory, D., Paule, L., Shvadchak, I. M., Popescu, F., Sulkowska, M., Hynek, V., & Longauer, R. (2003). Spatial patterns of the genetic differentiation in European beech (Fagus sylvatica L.) at allozyme loci in the Carpathians and the adjacent regions. Silvae Genetica, 52(2), 78–83.

Grinde, K. E., Browning, B. L., Reiner, A. P., Thornton, T. A., & Browning, S. R. (2024). Adjusting for principal components can induce spurious associations in genome-wide association studies in admixed populations. bioRxiv.

Guevara, M. Á., Sánchez-Gómez, D., Vélez, M. D., de María, N., Díaz, L. M., Ramírez-Valiente, J. A., … & Cervera, M. T. (2022). Epigenetic and Genetic Variability in Contrasting Latitudinal Fagus sylvatica L. Provenances. Forests, 13(12), 1971.

Guyot, G., Baret, F., & Jacquemoud, S. (1992). Imaging spectroscopy for vegetation studies (Vol. 2, pp. 145–165). Kluwer Academic Publishers: Norwell, MA, USA.

Hazler, K., Comps, B., Sugar, I., Melovski, L., Tashev, A., & Gracan, J. (1997). Genetic structure of Fagus sylvatica L. populations in Southeastern Europe. Silvae Genetica, 46(4), 229–235.

Hector, A., & Bagchi, R. (2007). Biodiversity and ecosystem multifunctionality. Nature, 448(7150), 188–190.

Hector, A., Hautier, Y., Saner, P., Wacker, L., Bagchi, R., Joshi, J., … & Loreau, M. (2010). General stabilizing effects of plant diversity on grassland productivity through population asynchrony and overyielding. Ecology, 91(8), 2213–2220.

Helfenstein, I. S., Schneider, F. D., Schaepman, M. E., & Morsdorf, F. (2022). Assessing biodiversity from space: Impact of spatial and spectral resolution on trait-based functional diversity. Remote Sensing of Environment, 275, 113024.

Higashi, Y., & Saito, K. (2019). Lipidomic studies of membrane glycerolipids in plant leaves under heat stress. Progress in lipid research, 75, 100990.

Hinney, A., Nguyen, T. T., Scherag, A., Friedel, S., Brönner, G., Müller, T. D., … & Hebebrand, J. (2007). Genome wide association (GWA) study for early onset extreme obesity supports the role of fat mass and obesity associated gene (FTO) variants. PloS one, 2(12), e1361.

Hoban, S., Archer, F. I., Bertola, L. D., Bragg, J. G., Breed, M. F., Bruford, M. W., … & Hunter, M. E. (2022). Global genetic diversity status and trends: towards a suite of Essential Biodiversity Variables (EBVs) for genetic composition. Biological Reviews, 97(4), 1511–1538.

Huang, M., Liu, X., Zhou, Y., Summers, R. M., & Zhang, Z. (2019). BLINK: a package for the next level of genome-wide association studies with both individuals and markers in the millions. Gigascience, 8(2), giy154.

Hughes, A. R., Inouye, B. D., Johnson, M. T., Underwood, N., & Vellend, M. (2008). Ecological consequences of genetic diversity. Ecology letters, 11(6), 609–623.

IPBES Intergovernmental Science-Policy Platform on Biodiversity and Ecosystem Services (IPBES). (2019). Global assessment report on biodiversity and ecosystem services of the Intergovernmental Science-Policy Platform on Biodiversity and Ecosystem Services.

Isbell, F., Calcagno, V., Hector, A., Connolly, J., Harpole, W. S., Reich, P. B., … & Loreau, M. (2011). High plant diversity is needed to maintain ecosystem services. Nature, 477(7363), 199–202.

Isbell, F., Craven, D., Connolly, J., Loreau, M., Schmid, B., Beierkuhnlein, C., … & Eisenhauer, N. (2015). Biodiversity increases the resistance of ecosystem productivity to climate extremes. Nature, 526(7574), 574–577.

Jacquemoud, S., & Ustin, S. (2019). Leaf optical properties. Cambridge University Press.

Jenkins, C. N., & Joppa, L. (2009). Expansion of the global terrestrial protected area system. Biological conservation, 142(10), 2166–2174.

Johnson, C. N., Balmford, A., Brook, B. W., Buettel, J. C., Galetti, M., Guangchun, L., & Wilmshurst, J. M. (2017). Biodiversity losses and conservation responses in the Anthropocene. Science, 356(6335), 270–275.

Johnston, R., Jones, K., & Manley, D. (2018). Confounding and collinearity in regression analysis: a cautionary tale and an alternative procedure, illustrated by studies of British voting behaviour. Quality & quantity, 52, 1957–1976.

Lazic, D., Geßner, C., Liepe, K. J., Lesur-Kupin, I., Mader, M., Blanc-Jolivet, C., … & Müller, N. A. (2024). Genomic variation of European beech reveals signals of local adaptation despite high levels of phenotypic plasticity. Nature Communications, 15(1), 8553.

Li, C., Czyż, E. A., Halitschke, R., Baldwin, I. T., Schaepman, M. E., & Schuman, M. C. (2023). Evaluating potential of leaf reflectance spectra to monitor plant genetic variation. Plant Methods, 19(1), 108.

Lou, R. N., Jacobs, A., Wilder, A. P., & Therkildsen, N. O. (2021). A beginner’s guide to low-coverage whole genome sequencing for population genomics. Molecular Ecology, 30(23), 5966–5993.

Madritch, M. D., Kingdon, C. C., Singh, A., Mock, K. E., Lindroth, R. L., & Townsend, P. A. (2014). Imaging spectroscopy links aspen genotype with below-ground processes at landscape scales. Philosophical Transactions of the Royal Society B: Biological Sciences, 369(1643), 20130194.

Magri, D., Vendramin, G. G., Comps, B., Dupanloup, I., Geburek, T., Gömöry, D., … & De Beaulieu, J. L. (2006). A new scenario for the Quaternary history of European beech populations: palaeobotanical evidence and genetic consequences. New phytologist, 171(1), 199–221.

Maruki, T., & Lynch, M. (2015). Genotype-frequency estimation from high-throughput sequencing data. Genetics, 201(2), 473–486.

Maruki, T., & Lynch, M. (2017). Genotype calling from population-genomic sequencing data. G3: Genes, Genomes, Genetics, 7(5), 1393–1404.

Meger, J., Ulaszewski, B., & Burczyk, J. (2021). Genomic signatures of natural selection at phenology-related genes in a widely distributed tree species Fagus sylvatica L. BMC genomics, 22(1), 1–20.

Meireles, J. E., Cavender-Bares, J., Townsend, P. A., Ustin, S., Gamon, J. A., Schweiger, A. K., … & O’Meara, B. C. (2020). Leaf reflectance spectra capture the evolutionary history of seed plants. New Phytologist, 228(2), 485–493.

Miller, J. R., Steven, M. D., & Demetriades-Shah, T. H. (1992). Reflection of layered bean leaves over different soil backgrounds: measured and simulated spectra. International Journal of Remote Sensing, 13(17), 3273–3286.

Mishra, B., Ulaszewski, B., Meger, J., Aury, J. M., Bodénès, C., Lesur-Kupin, I., … & Thines, M. (2022). A Chromosome-level genome assembly of the European Beech (Fagus sylvatica) reveals anomalies for organelle DNA integration, repeat Content and distribution of SNPs. Frontiers in genetics, 12, 691058.

Mora, C., Andréfouët, S., Costello, M. J., Kranenburg, C., Rollo, A., Veron, J., … & Myers, R. A. (2006). Coral reefs and the global network of marine protected areas. Science, 312(5781), 1750–1751.

Østerberg, J. T., & Palmgren, M. (2018). Heavy metal pumps in plants: structure, function and origin. Advances in Botanical Research, 87, 57–89.

Perrings, C., Duraiappah, A., Larigauderie, A., & Mooney, H. (2011). The biodiversity and ecosystem services science-policy interface. Science, 331(6021), 1139–1140.

Petit, R. J., Aguinagalde, I., de Beaulieu, J. L., Bittkau, C., Brewer, S., Cheddadi, R., … & Vendramin, G. G. (2003). Glacial refugia: hotspots but not melting pots of genetic diversity. science, 300(5625), 1563–1565.

Pfenninger, M., Reuss, F., Kiebler, A., Schönnenbeck, P., Caliendo, C., Gerber, S., … & Feldmeyer, B. (2020). Genomic basis of drought resistance in Fagus sylvatica. bioRxiv, 2020-12.

Pluess, A. R., & Weber, P. (2012). Drought-adaptation potential in Fagus sylvatica: linking moisture availability with genetic diversity and dendrochronology. PloS one, 7(3), e33636.

Podani, J., & Miklós, I. (2002). Resemblance coefficients and the horseshoe effect in principal coordinates analysis. Ecology, 83(12), 3331–3343.

Purvis, A., & Hector, A. (2000). Getting the measure of biodiversity. Nature, 405(6783), 212–219.

Reich, P. B., Tilman, D., Isbell, F., Mueller, K., Hobbie, S. E., Flynn, D. F., & Eisenhauer, N. (2012). Impacts of biodiversity loss escalate through time as redundancy fades. Science, 336(6081), 589–592.

Schaepman, M. E., Jehle, M., Hueni, A., D’Odorico, P., Damm, A., Weyermann, J., … & Itten, K. I. (2015). Advanced radiometry measurements and Earth science applications with the Airborne Prism Experiment (APEX). Remote Sensing of Environment, 158, 207–219.

Schaepman, M. E., Ustin, S. L., Plaza, A. J., Painter, T. H., Verrelst, J., & Liang, S. (2009). Earth system science related imaging spectroscopy—An assessment. Remote Sensing of Environment, 113, S123–S137.

Scholes, R. J., Mace, G. M., Turner, W., Geller, G. N., Jürgens, N., Larigauderie, A., … & Mooney, H. A. (2008). Toward a global biodiversity observing system. Science, 321(5892), 1044–1045.

Scott, J. M., Csuti, B., Estes, J. E., & Anderson, H. (1989). Status assessment of biodiversity protection. Conservation Biology, 3(1), 85–87.

Seeley, M. M., Martin, R. E., Vaughn, N. R., Thompson, D. R., Dai, J., & Asner, G. P. (2023). Quantifying the variation in reflectance spectra of Metrosideros polymorpha canopies across environmental gradients. Remote Sensing, 15(6), 1614.

Stefanini, C., Csilléry, K., Ulaszewski, B., Burczyk, J., Schaepman, M. E., & Schuman, M. C. (2023). A novel synthesis of two decades of microsatellite studies on European beech reveals decreasing genetic diversity from glacial refugia. Tree Genetics & Genomes, 19(1), 3.

Sutherland, W. J., Pullin, A. S., Dolman, P. M., & Knight, T. M. (2004). The need for evidence-based conservation. Trends in ecology & evolution, 19(6), 305–308.

Tang, T., Zhang, N., Bongers, F. J., Staab, M., Schuldt, A., Fornoff, F., … & Liu, X. (2022). Tree species and genetic diversity increase productivity via functional diversity and trophic feedbacks. Elife, 11, e78703.

Thiébaut, B., Lumaret, R., & Vernet, P. H. (1982). The bud enzymes of beech (Fagus sylvatica L.) genetic distinction and analysis of polymorphism in several French populations.

Tilman, D., & Downing, J. A. (1994). Biodiversity and stability in grasslands. Nature, 367(6461), 363–365.

Turner, W. (2014). Sensing biodiversity. Science, 346(6207), 301–302.

van Moorsel, S. J., Hahl, T., Petchey, O. L., Ebeling, A., Eisenhauer, N., Schmid, B., & Wagg, C. (2021). Co- occurrence history increases ecosystem stability and resilience in experimental plant communities. Ecology, 102(1), e03205.

Wang, J., & Zhang, Z. (2021). GAPIT version 3: boosting power and accuracy for genomic association and prediction. Genomics, Proteomics and Bioinformatics, 19(4), 629–640.

Wilson, E. O. (1988). Biodiversity.

Yamasaki, E., Altermatt, F., Cavender-Bares, J., Schuman, M. C., Zuppinger-Dingley, D., Garonna, I., … & Shimizu, K. K. (2017). Genomics meets remote sensing in global change studies: monitoring and predicting phenology, evolution and biodiversity. Current opinion in environmental sustainability, 29, 177–186.

Zhang, P., Wang, F., Zhang, L. F., Rui, Q., & Xu, L. L. (2006). The role of serine endopeptidase in cucumber leaf senescence. Zhi wu Sheng li yu fen zi Sheng wu xue xue bao= Journal of Plant Physiology and Molecular Biology, 32(5), 593–599.

Zheng, Z., Zeng, Y., Schuman, M. C., Jiang, H., Schmid, B., Schaepman, M. E., & Morsdorf, F. (2022). Individual tree-based vs pixel-based approaches to mapping forest functional traits and diversity by remote sensing. International Journal of Applied Earth Observation and Geoinformation, 114, 103074.

