## Supporting information for "Associations of genetic and spectral diversity in European beech"

Article title: Associations of genetic and spectral diversity in European beech

The following Supporting Information is available for this article:

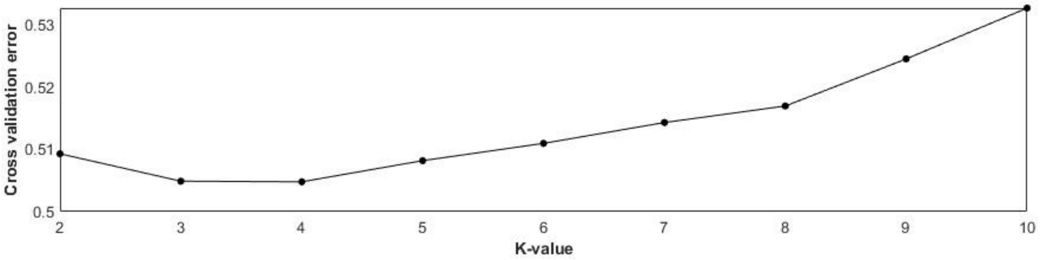


**Fig. S1** Plot of *ADMIXTURE* cross validation error from K=1 through K=10. The number of genetic clusters obtained using the SNPs data was set to 4 based on the smallest cross-validation error.

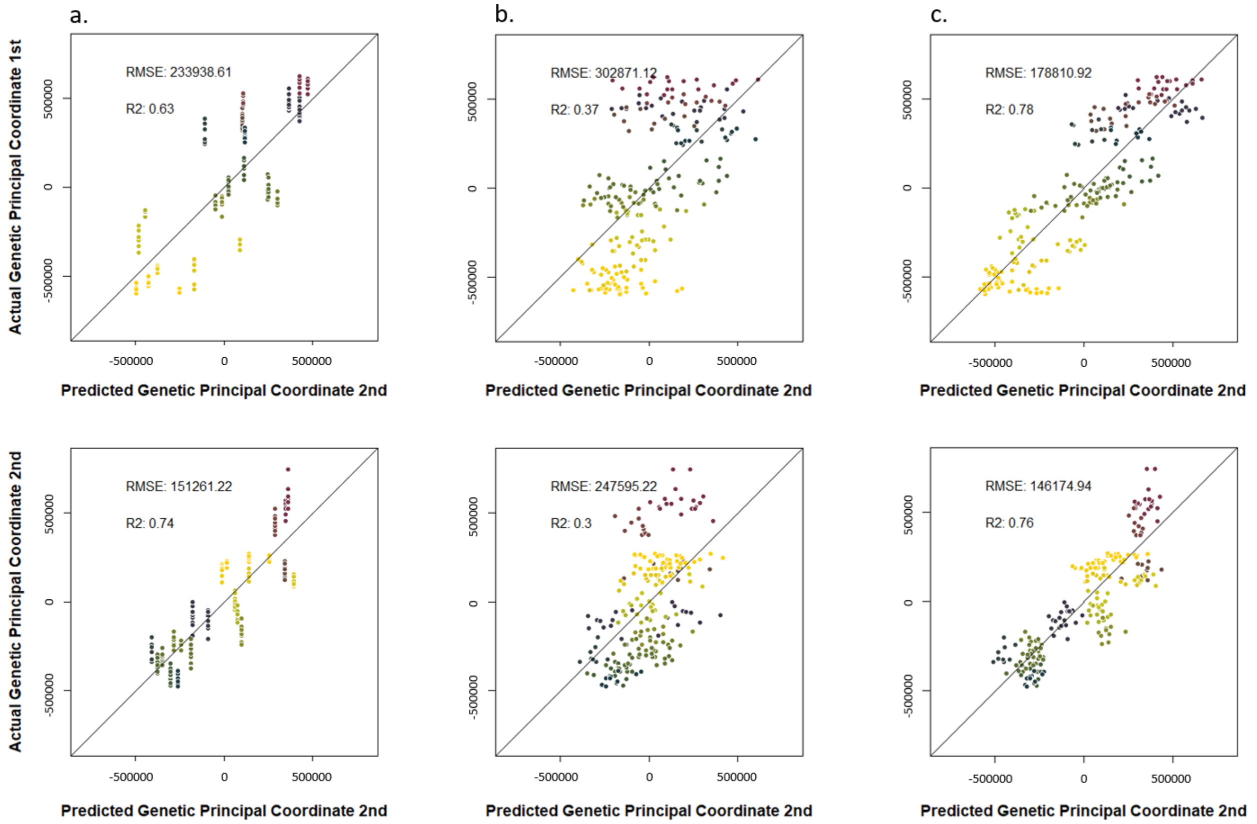


**Fig. S2** RDA models prediction of genetic structure along the first and second genetic PCOs in comparison to the actual PCOs based on genetic distances between sampled *F. sylvatica* individuals derived from pairwise allele counts. The RDA models were build using environmental information (**a**), spectral information (**b**) and combined environmental with spectral information (**c**).

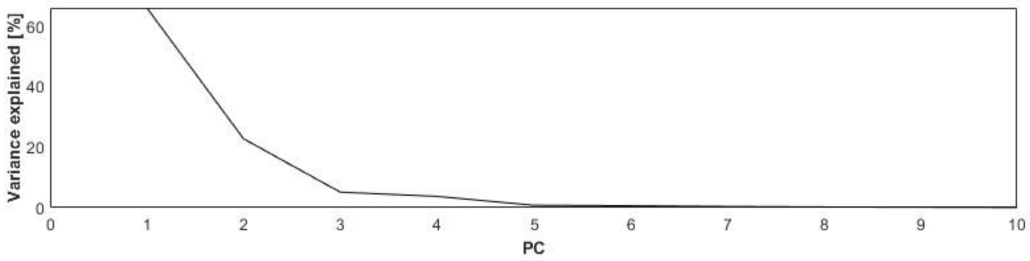


**Fig. S3** Variance in reflectance spectra explained by Principal Components (216 spectral variables measured on n = 219 individuals).

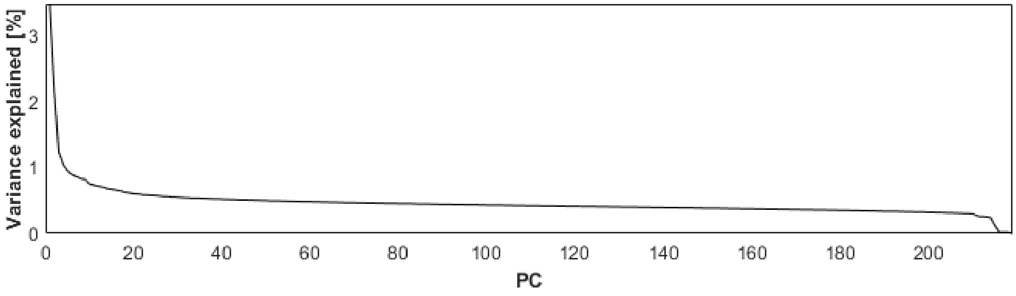


**Fig. S4** Variance in genomic structure explained by Principal Components (n = 219 individuals).

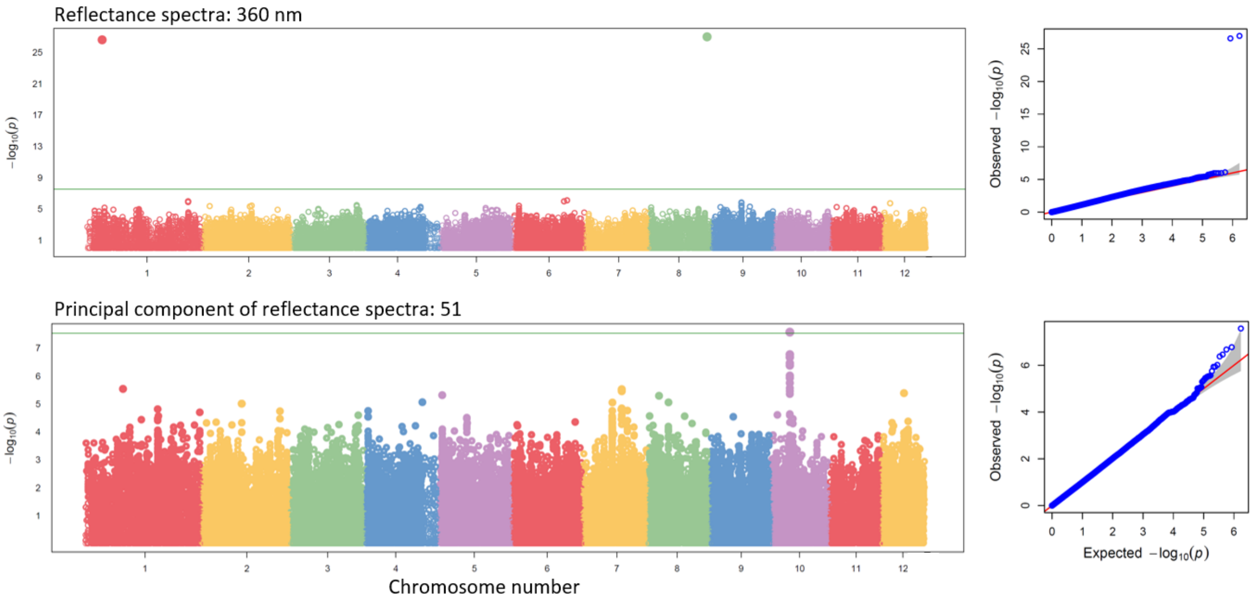


**Fig. S7** Manhattan and QQ plots for two spectral variables for which associations with SNPs variation was significant according to Bonferroni criterium that accounts on the multiple statistical tests to reduce false positives and which were associated with SNPs located within annotated genes of *F. sylvatica*.

**Tab. S1** Locations of the sites and the dates of the foliage collection for spectral and genomic analyses.

| Site ID | Country | Collection date | Longitude | Latitude |
|---|---|---|---|---|
| BES | Belgium | 2020/09/02 | 8.372298 | 47.475272 |
| CHL | Switzerland | 2020/08/07 | 4.422831 | 50.750554 |
| ESM | Spain | 2020/09/23&24 | -1.810131 | 41.790907 |
| ESP | Spain | 2020/09/24 | -0.673556 | 42.509507 |
| FRB | France | 2020/09/19 | 5.769441 | 43.328252 |
| FRC | France | 2020/08/23 | 9.162245 | 41.882290 |
| FRM | France | 2020/08/25 | 3.031944 | 42.491384 |
| FRV | France | 2020/08/22 | 9.097174 | 42.114789 |
| HRP | Croatia | 2020/09/16 | 15.467072 | 44.360311 |
| ITB | Italy | 2020/08/11&21 | 12.156174 | 42.173892 |
| ITE | Italy | 2020/08/17 | 15.044973 | 37.708737 |
| ITF | Italy | 2020/08/14 | 15.753308 | 40.505757 |
| ITM | Italy | 2020/08/16 | 14.041612 | 37.857776 |
| ITP | Italy | 2020/08/19 | 16.109641 | 39.905240 |
| ITV | Italy | 2020/08/13 | 15.631165 | 40.948701 |
| PLB | Poland | 2020/09/11 | 22.472191 | 49.232498 |
| PLM | Poland | 2020/09/07 | 21.471417 | 53.713383 |
| PLW | Poland | 2020/09/09 | 18.860433 | 49.669433 |
| RSB | Serbia | 2020/09/14 | 13.218648 | 40.072190 |
| RSF | Serbia | 2020/09/13 | 19.635920 | 45.136552 |
| RSO | Romania | 2021/07/31 | 25.166683 | 45.688390 |
| SEB | Sweden | 2020/09/04 | 13.208720 | 55.560152 |
| SLK | Slovenia | 2020/09/17 | 14.763337 | 45.542632 |
